## Supplementary material for "Functional recruitment and connectivity of the cerebellum supports the emergence of Theory of Mind in early childhood"

**Supplementary figure 1**

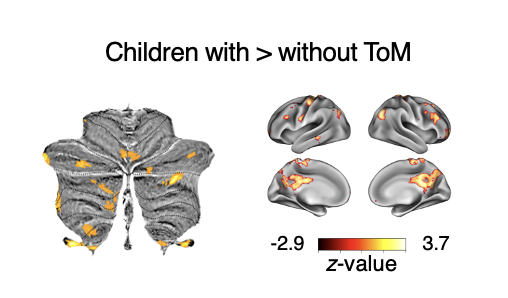

**Supplementary figure 1. Functional activation controlling for IQ.** General linear model (GLM) of activation differences between children with and without ToM with children’s ToM task performance (0-6) and IQ score (WPPSI or KBIT-II) as a predictor (FDR-corrected on a *q* = .05 level).

**Supplementary figure 2**

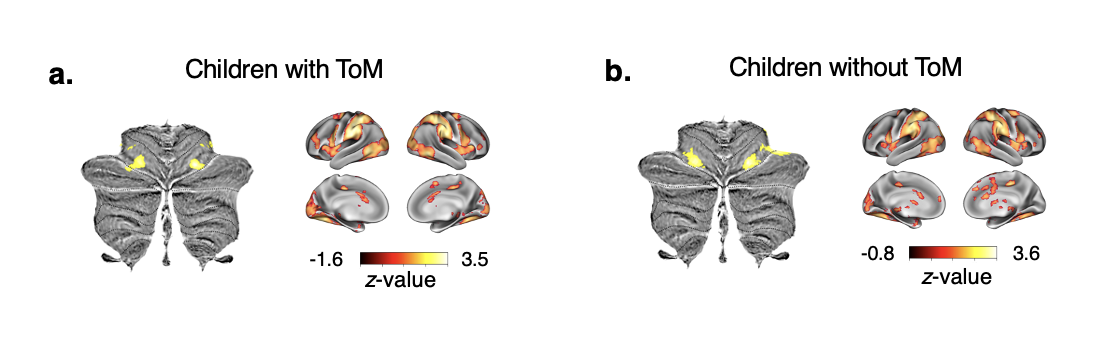

**Supplementary figure 2. Functional activation during movie scenes depicting bodily transformation (pain) in children with and without ToM abilities** (FDR-corrected on a *q* = .05 level). **a.** One-sample *t*-test showing activations for bodily transformation vs. ToM movie scenes in children with ToM abilities. **b.** One-sample *t*-test showing activations for bodily transformation vs. ToM movie scenes in children without ToM abilities.

**Supplementary figure 3**

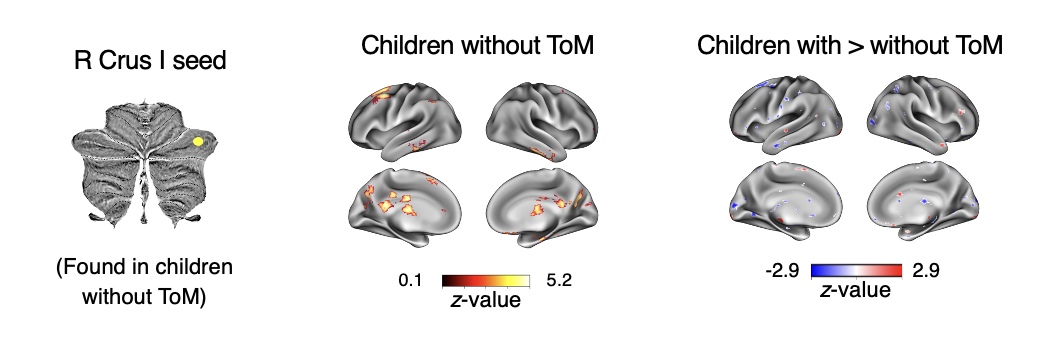

**Supplementary figure 3. Seed-to-voxel functional connectivity between cerebellar ToM clusters and the cerebral cortex in children without ToM abilities** (FDR-corrected on a *q* = .05 level). Left: One-sample *t*-test of connectivity of rCrus I (cluster found in children without ToM abilities) with the cerebral cortex in children without ToM abilities. Right: Two-sample *t*-test of connectivity differences of rCrus I between children with and without ToM abilities.

**Supplementary figure 4**

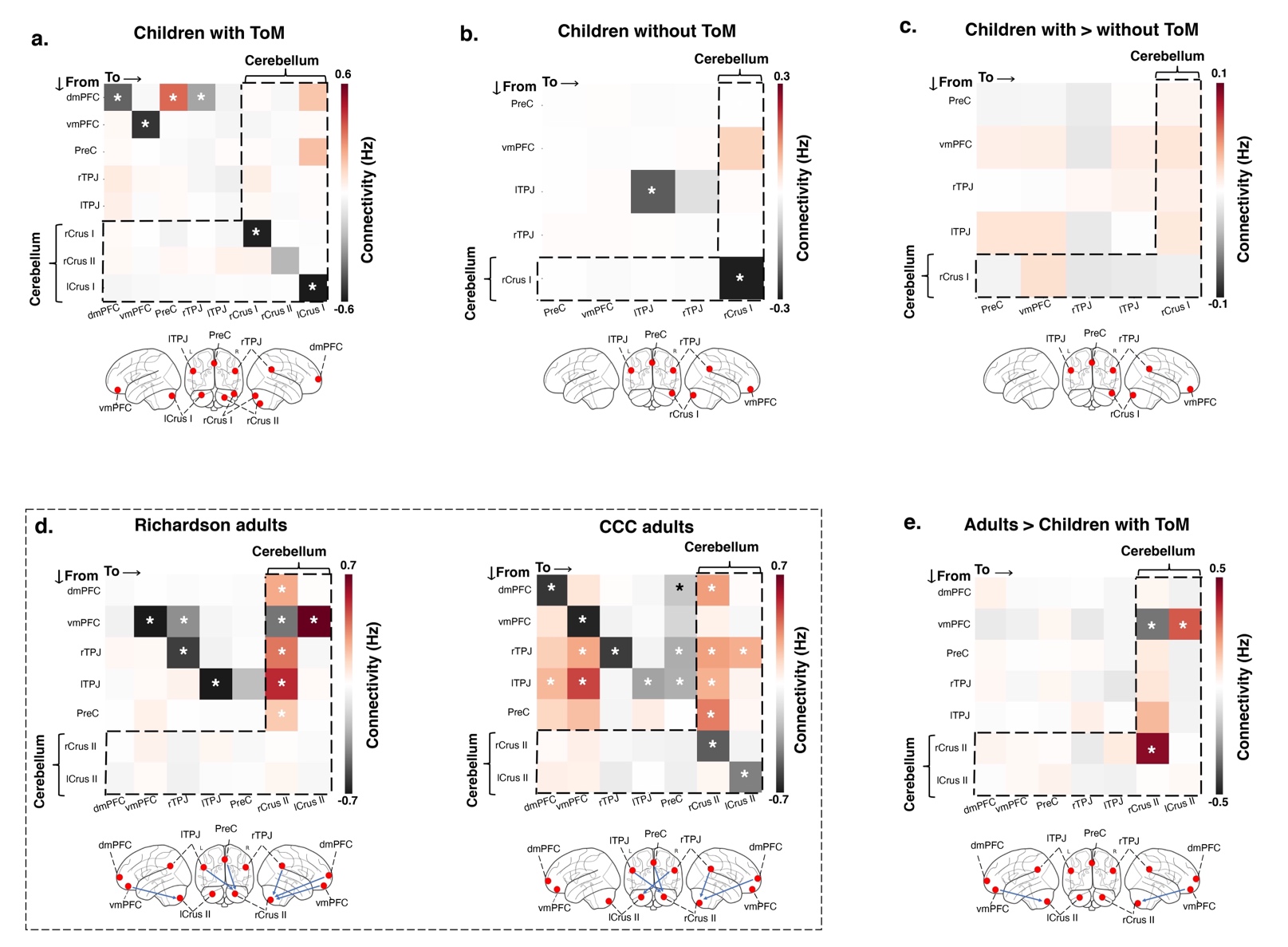

**Supplementary figure 4. Dynamic causal modeling (DCM) of the cerebellum and the cerebral ToM network in children with ToM abilities and adults.** Averaged modulatory (task-dependent) connections in children with and without ToM abilities and adults (in units of Hz). The vertical axis represents connections that originate from a seed region and terminate to a target region (represented in the horizontal axis). **a, b.** Modulatory connections in children (a) with and (b) without ToM abilities, using ToM activation clusters (identified via the in-scanner ToM task) as ROIs. **c.** Connectivity differences between children with and without ToM abilities, identified by adding group type (ToM abilities, no ToM abilities) as a covariate in the model. **d.** Modulatory connections in adults in the Richardson et al. (2018) and CCC samples using ToM ROIs identified in a functional atlas of ToM activations in adults (King et al., 2019). **e.** Connectivity differences between children with ToM abilities and adults (in the Richardson et al., 2018 sample), using group type (adult, child) as a covariate. Green arrows in the glass brains represent connections from the cerebellum to the cerebral cortex. Blue arrows represent connections from the cerebral cortex to the cerebellum. * = Bayesian posterior probability > .95. Abbreviations: dmPFC = dorsomedial prefrontal cortex; vmPFC = ventromedial prefrontal cortex; PreC = precuneus; r/lTPJ = right/left temporoparietal junction.

**Supplementary figure 5**

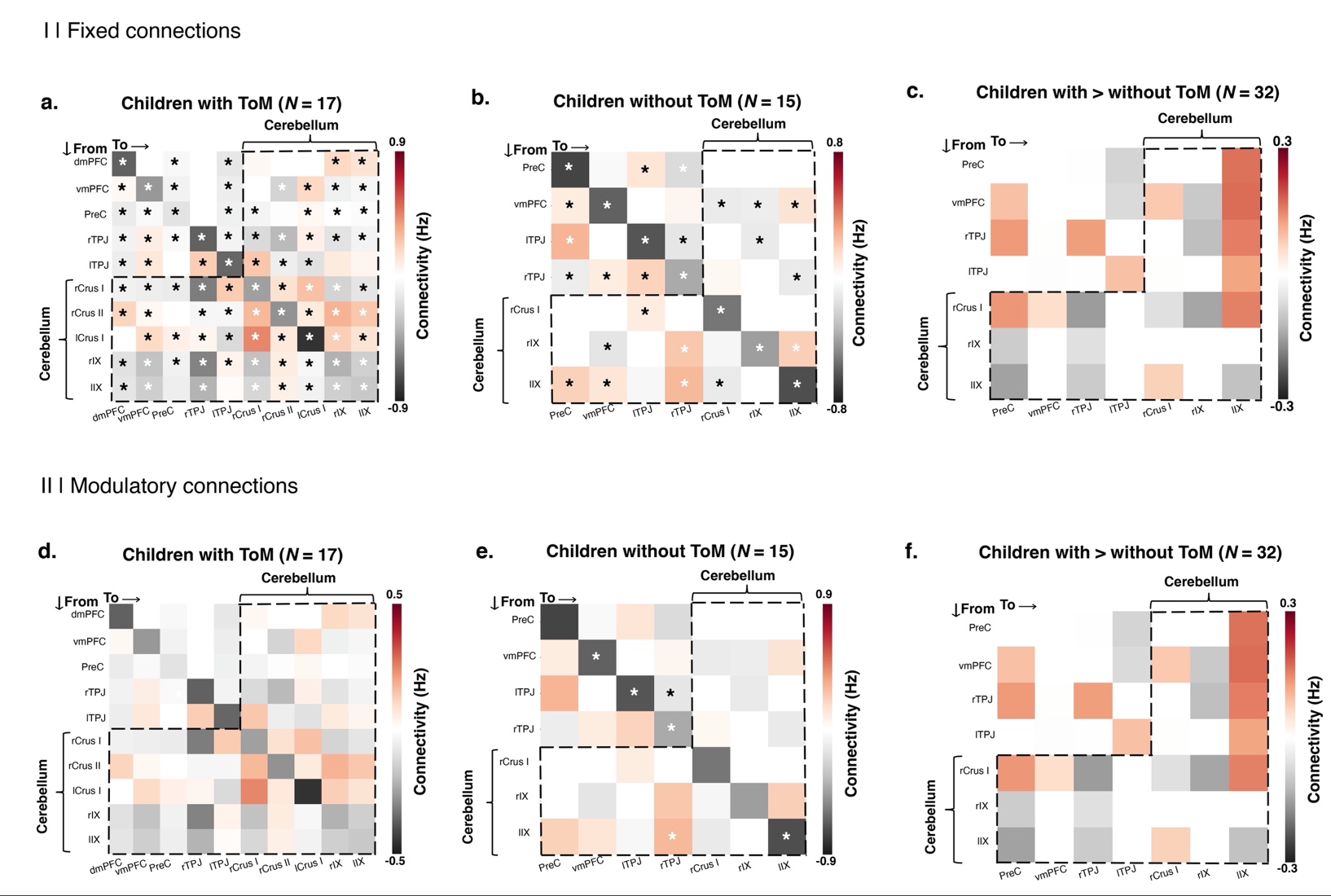

**Supplementary figure 5. Dynamic causal modeling (DCM) of the cerebellum and the cerebral ToM network in a subset of children with activations in the bilateral IX.** Averaged fixed (task-independent; I) and modulatory (task-dependent; II) connections in children with and without ToM abilities and adults (in units of Hz). The vertical axis represents connections that originate from a seed region and terminate to a target region (represented in the horizontal axis). **a, b, d, e.** Modulatory connections in children (a, d) with and (b, e) without ToM abilities, using ToM activation clusters (identified via the in-scanner ToM task) as ROIs. **c, f.** Connectivity differences between children with and without ToM abilities, identified by adding group type (ToM abilities, no ToM abilities) as a covariate in the model. * = Bayesian posterior probability > .95. Abbreviations: dmPFC = dorsomedial prefrontal cortex; vmPFC = ventromedial prefrontal cortex; PreC = precuneus; r/lTPJ = right/left temporoparietal junction.

**Supplementary table 1**

*MNI coordinates of ToM ROIs in all children*

| **ROI** | **MNI Coordinates** | | |
| --- | --- | --- | --- |
|  | **x** | **y** | **z** |
| *Cerebral cortex* |  |  |  |
| vmPFC | 4 | 51 | -10 |
| PreC | 0 | -46 | 36 |
| rTPJ | 43 | -65 | 38 |
| lTPJ | -48 | -65 | 38 |
| *Cerebellum* |  |  |  |
| rCrus I | 52 | -68 | -26 |
| rIX | 2 | -46 | -44 |
| lIX | -8 | -46 | -44 |

Abbreviations: vmPFC = ventromedial prefrontal cortex; PreC = precuneus; r/lTPJ = right/left temporoparietal junction.

**Supplementary table 2**

*MNI coordinates of ToM ROIs in ToM passers*

| **ROI** | **MNI Coordinates** | | |
| --- | --- | --- | --- |
|  | **x** | **y** | **z** |
| *Cerebral cortex* |  |  |  |
| dmPFC | -6 | 45 | 42 |
| vmPFC | 4 | 56 | 0 |
| PreC | 14 | -48 | 30 |
| rTPJ | 42 | -64 | 40 |
| lTPJ | -48 | -67 | 37 |
| *Cerebellum* |  |  |  |
| rCrus I | 53 | -56 | -30 |
| lCrus I | -30 | -84 | -30 |
| rCrus II | 22 | -87 | -36 |
| rIX | 8 | -50 | -44 |
| lIX | -8 | -52 | -38 |

Abbreviations: dmPFC = dorsomedial prefrontal cortex; vmPFC = ventromedial prefrontal cortex; PreC = precuneus; r/lTPJ = right/left temporoparietal junction.

**Supplementary table 3**

*MNI coordinates of ToM ROIs in ToM non-passers*

| **ROI** | **MNI Coordinates** | | |
| --- | --- | --- | --- |
|  | **x** | **y** | **z** |
| *Cerebral cortex* |  |  |  |
| vmPFC | 2 | 55 | -11 |
| PreC | 14 | -55 | 26 |
| rTPJ | 42 | -67 | 38 |
| lTPJ | -34 | -72 | 43 |
| *Cerebellum* |  |  |  |
| rCrus I | 54 | -68 | -30 |
| rIX | 3 | -44 | -44 |
| lIX | -8 | -44 | -44 |

Abbreviations: vmPFC = ventromedial prefrontal cortex; PreC = precuneus; r/lTPJ = right/left temporoparietal junction.

**Supplementary table 4**

*Number of participants with ToM ROIs defined at p > .05*

| **Group** |  | **rTPJ** | | **lTPJ** | | **dmPFC** | | **vmPFC** | | **PreC** | | **rCrus II** | | **lCrus II** | | **rCrus I** | | **lCrus I** | |
| --- | --- | --- | --- | --- | --- | --- | --- | --- | --- | --- | --- | --- | --- | --- | --- | --- | --- | --- | --- |
|  |  | ***p*<.1** | ***p*<1** | ***p*<.1** | ***p*<1** | ***p*<.1** | ***p* <1** | ***p*<.1** | ***p*<1** | ***p*<.1** | ***p*<1** | ***p*<.1** | ***p*<1** | ***p*<.1** | ***p*<1** | ***p*<.1** | ***p*<1** | ***p*<.1** | ***p*<1** |
| *Adults* | ***N*** |  |  |  |  |  |  |  |  |  |  |  |  |  |  |  |  |  |  |
| Richardson et al. (2018) | 22 | 0 | 0 | 0 | 0 | 1 | 0 | 1 | 1 | 0 | 1 | 1 | 0 | 1 | 0 | - | - | - | - |
| CCC | 56 | 0 | 0 | 0 | 0 | 2 | 0 | 1 | 1 | 0 | 0 | 2 | 3 | 0 | 3 | - | - | - | - |
| *Children* | ***N*** |  |  |  |  |  |  |  |  |  |  |  |  |  |  |  |  |  |  |
| All | 41 | 0 | 0 | 1 | 0 | - | - | 3 | 1 | 3 | 0 | - | - | - | - | 3 | 9 | - | - |
| ToM passers | 22 | 0 | 0 | 0 | 0 | 1 | 0 | 0 | 0 | 0 | 0 | 7 | 1 | - | - | 4 | 4 | 3 | 4 |
| ToM non-passers | 19 | 0 | 0 | 1 | 0 | - | - | 3 | 1 | 2 | 0 | - | - | - | - | 3 | 9 | - | - |

Dashes (-) represent ROIs that were not applicable in a given sample. *N* refers to the total number of subjects in a given sample. Abbreviations: dmPFC = dorsomedial prefrontal cortex; vmPFC = ventromedial prefrontal cortex; PreC = precuneus; r/lTPJ = right/left temporoparietal junction.

**Supplementary methods**

***ToM specificity.*** We performed additional analyses to ensure that the observed differences in functional activations between ToM passers and non-passers (**Figure 1**) were specific to children’s ToM abilities and not driven by the development of general cognitive abilities. First, we ran a second-level general linear model (GLM) on the entire developmental sample, including children’s ToM score (ratio predictor: 0-6) as well as children’s IQ. Given that children performed a different out-of-scanner IQ test depending on their age (see **Behavioral task battery**), we added two IQ predictors in our model: *IQ test type,* a binary categorical variable indicating which IQ test a child completed (WPPSI or KBIT-II), and *IQ score*, a continuous variable including the pooled standardized scores of the two IQ tests. The effect of ToM score on functional activation in the cerebellum (**Figure 1b**) remained similar in bilateral medial Crus I (rCrus I: 19 -85 -30; lCrus I: -40 -75 -35) and Crus II (rCrus II: 42 -74 -45; lCrus II: -12 -69 -43) (*p_uncorr._* < . 001, FDR-corrected: *q* = .05) (**Supplementary figure 1**). It should be noted that IQ score was moderately correlated with ToM score (Spearman’s *r*(39) = .72, *p* < .001).

Additionally, we examined if functional activation in the cerebellum for bodily transformations (i.e., pain) was also affected by children’s out-of-scanner ToM task performance. This would mean that the functional differences between ToM passers and non-passers were not necessarily driven by ToM abilities, but some other process (e.g., simply reflected changes due to age). On a single-subject level, we followed the method described in **Contrast analyses**, but focused on the pain > ToM contrast (as opposed to the ToM > pain contrast). On the group level, we performed two separate one-sample *t*-tests in the groups of ToM passers and non-passers (*p_uncorr._* < . 001, FDR-corrected: *q* = .05). We observed functional clusters in the bilateral IV that did not overlap with the clusters found for the ToM > pain contrast **(Figure 1c-d**), both in ToM passers (rIV: 25 -71 -23; lIV: -34 -73 -24) and non-passers (rIV: 26 -74 -22; lIV: -29 -74 -22), further ensuring the ToM specificity of our results (**Supplementary figure 2**).
